# Native-like SARS-CoV-2 spike glycoprotein expressed by ChAdOx1 nCoV-19/AZD1222 vaccine

**DOI:** 10.1101/2021.01.15.426463

**Authors:** Yasunori Watanabe, Luiza Mendonça, Elizabeth R. Allen, Andrew Howe, Mercede Lee, Joel D. Allen, Himanshi Chawla, David Pulido, Francesca Donnellan, Hannah Davies, Marta Ulaszewska, Sandra Belij-Rammerstorfer, Susan Morris, Anna-Sophia Krebs, Wanwisa Dejnirattisai, Juthathip Mongkolsapaya, Piyada Supasa, Gavin R. Screaton, Catherine M. Green, Teresa Lambe, Peijun Zhang, Sarah C. Gilbert, Max Crispin

## Abstract

Vaccine development against the SARS-CoV-2 virus focuses on the principal target of the neutralizing immune response, the spike (S) glycoprotein. Adenovirus-vectored vaccines offer an effective platform for the delivery of viral antigen, but it is important for the generation of neutralizing antibodies that they produce appropriately processed and assembled viral antigen that mimics that observed on the SARS-CoV-2 virus. Here, we describe the structure, conformation and glycosylation of the S protein derived from the adenovirus-vectored ChAdOx1 nCoV-19/AZD1222 vaccine. We demonstrate native-like post-translational processing and assembly, and reveal the expression of S proteins on the surface of cells adopting the trimeric prefusion conformation. The data presented here confirms the use of ChAdOx1 adenovirus vectors as a leading platform technology for SARS-CoV-2 vaccines.

## Introduction

Vaccines against severe acute respiratory syndrome coronavirus-2 (SARS-CoV-2) are an essential countermeasure to stem the COVID-19 pandemic. Vaccine development efforts aim to produce both a strong T cell response and a neutralizing immune response against the virus, the main target being the spike (S) proteins that protrude from the viral envelope (*1*). The S protein is responsible for mediating host-cell entry, with the S1 and S2 subunits facilitating angiotensin-converting enzyme 2 (ACE2) receptor binding and membrane fusion, respectively (*2*–*4*). The SARS-CoV-2 S gene encodes the extensively glycosylated trimeric class I viral fusion protein with 22 N-linked glycans per protomer (*5*).

Whilst the coronavirus S glycoprotein is the principal target for SARS-CoV-2 vaccine design, leading vaccine candidates and recently licensed vaccines utilise a variety of constructs and strategies (*6*). For example, both Moderna’s mRNA-1273 and Pfizer’s BNT162b2 (*7*) encodes full length S with two mutations to stabilise the prefusion conformation (*8*), and Sinovac’s CoronaVac inactivated virus vaccine presents the wild-type S on the viral surface (*9*), although the majority of spikes are in the postfusion conformation (*10*). One key aim for SARS-CoV-2 vaccine development is to elicit a robust immune response against the spike, and more specifically the RBD, where many neutralizing epitopes are located (*10*–*15*). To this end, many vaccine candidates include (two or more) stabilising mutations in the S protein, such that the protein maintains the prefusion conformation and avoids shedding of S1 (*3*).

The replication-deficient chimpanzee adenovirus-vectored vaccine, ChAdOx1 nCoV-19/AZD1222 (hereafter referred to as ChAdOx1 nCoV-19), encodes the full-length wild-type SARS-CoV-2 spike protein. ChAdOx1 nCoV-19 has previously been shown to elicit not only strong neutralizing antibody responses, but also robust spike-specific T-cell responses (*16*–*19*). Although adenovirus-vectored vaccines are a promising way to deliver viral glycoprotein antigens, the processing and presentation of the SARS-CoV-2 spike is yet to be characterised. Understanding the molecular features of the expressed viral antigen is important for the interpretation of the immune response to this vaccine. Here, we determine the native-like functionality, prefusion trimeric structure and glycosylation of SARS-CoV-2 S protein expressed from the ChAdOx1 nCoV-19 vaccine.

## Results and Discussion

### Expression of prefusion conformation SARS-CoV-2 S glycoprotein on cell surfaces upon ChAdOx1 nCoV-19 infection

ChAdOx1 nCoV-19 encodes a wild-type S sequence, including the transmembrane domains, which is trafficked to the cell surface with a tissue plasminogen activator (tPA) signal sequence (**Fig. 1A**). Using flow cytometry, we first detected the presence of the S glycoprotein at the cell surface of ChAdOx1 nCoV-19 infected HeLa S3 cells (**Fig. 1B-E**). Sera from mice vaccinated with ChAdOx1 nCoV-19 was used to detect the expression level of S at the cell surface, revealing ~60-70% of all cells expressing S (range of duplicate averages across three experimental repeats) (**Sup. Fig. 1**). A ChAdOx1 vaccine encoding an irrelevant filovirus antigen (EBOV) was used as a negative control (*20*), which showed a low level of binding to the anti-ChAdOx1 nCoV-19 vaccine serum (~0.5-2% cells across all experiments). This observation accounts for antibodies raised against the vector itself rather that the vaccine antigen. Subsequently, we sought to examine the properties of the cell surface expressed S protein using recombinant ACE2 and a panel of human mAbs which bind to specific regions of S **(Fig. 1B)**. Binding of infected cells to recombinant ACE2 confirms the correct folding of S and native presentation and functionality of the RBD (**Fig. 1B**). This observation is further supported by the binding of the human mAbs, in particular Ab45 which recognises RBD, Ab71 which recognises the trimeric spike and Ab111 which recognises the NTD. Ab44 which recognises S2 also demonstrates considerable binding. These data confirm significant presence of the prefusion trimer at the cell surface. In the absence of a postfusion specific anti-S2 antibody we were unable to quantify if some postfusion spike is present at the cell surface.

**Figure 1.**
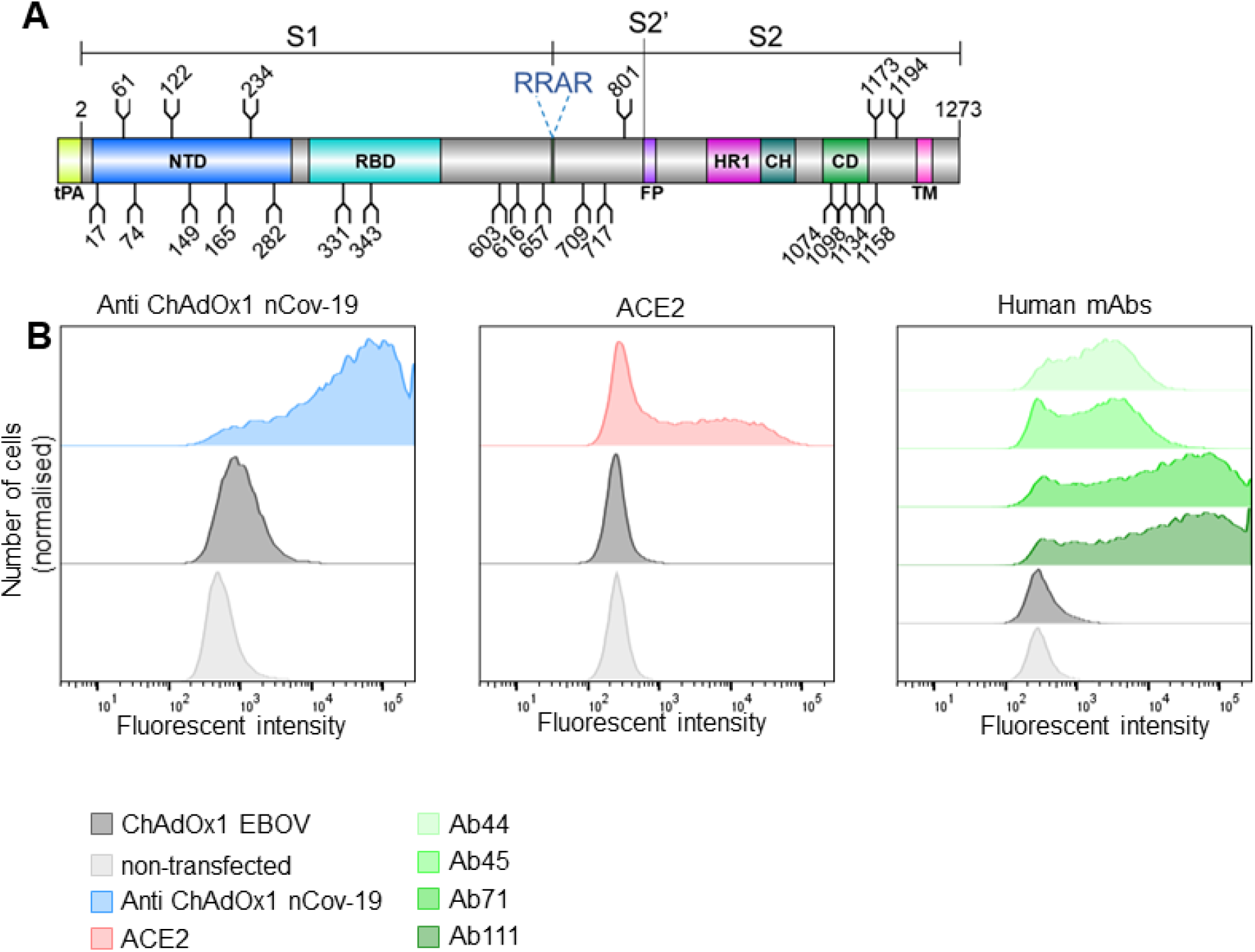
ChAdOx1 nCoV-19 produces membrane associated SARS-CoV-2 S glycoprotein in native conformations able to bind its host receptor, ACE2. **(A)** Schematic representation of the vaccine encoded SARS-CoV-2 S protein, showing the position of N-linked glycosylation amino-acid sequons (NXS/T, where X≠P) as branches. Protein domains are illustrated: N-terminal domain (NTD), receptor-binding domain (RBD), fusion peptide (FP), heptad repeat 1 (HR1), central helix (CH), connector domain (CD), and transmembrane domain (TM), with the additional tPA secretion signal at the N-terminus. **(B)** HeLa S3 cells were infected with ChAdOx1 nCoV-19 and incubated with either recombinant ACE2, anti-ChAdOx1 nCoV-19 (derived from vaccinated mice) or a panel of human mAbs (Ab44, Ab45, Ab71 and Ab111, which recognise, S2, RBD, trimeric S and NTD respectively) and compared to non-infected controls, analysed by flow cytometry. (Left). Relative frequency of cells and AlexaFluor 488 fluorescence associated with anti-spike detection is plotted. Left, (blue) anti-ChAdOx1 nCoV-19, middle (red), ACE2 and right (shades of green) human mAbs. In dark grey cells infected with an irrelevant ChAdOx1 vaccine and in light grey non-infected cells are shown as a control. Experimental replicates were performed two times and representative data shown.

Whilst this observation that the majority of cells infected with ChAdOx1 nCoV-19 present native-like spikes on the cell surface (**Fig. 1B**), it is interesting to note that a population may shed the S1 subunit. Whether this is a beneficial or detrimental feature with respect to the elicitation of immune responses during vaccination is unknown. Shedding of S1 subunits from viruses occurs during native infection (*21*–*24*), and the ChAdOx1 nCoV-19-derived S proteins mimics this native feature of the viral spike. Nevertheless it has been shown that vaccination with ChAdOx nCoV-19 elicits robust antibody and T cell responses (*16*–*19*).

### Structural analysis of membrane-associated ChAdOx1 nCoV-19 derived SARS-CoV-2 S protein

After the surface expression of the S protein driven by ChAdOx1 nCoV-19 vector infection was confirmed by cytometry, we sought to probe the structure of S proteins on native cell surfaces using cryo-electron tomography and subtomogram averaging (cryoET STA). We imaged U2OS and HeLa cells infected with ChAdOx1 nCoV-19. These cells were chosen for the thin cell peripheries which make them accessible for cryoET analysis. The tomograms revealed that surface of the cells is densely covered with protruding densities consistent with the size and shape to the prefusion conformation of SARS-CoV-2 S protein (**Fig. 2A & 2B, movie 1**). These densities are absent in control uninfected cells (**Sup. Fig. 2**). To determine whether these spikes represent the SARS-CoV-2 prefusion spike, we performed subtomogram averaging of 11391 spikes from cell surfaces using emClarity (*25*). The averaged density map, with three-fold symmetry applied, is at 11.6 Å resolution (at 0.5 FSC cut-off) (**Fig. 2C**), resolving the overall spike structure, which overlaps very well with prefusion spike atomic models in the literature (*4*, *26*–*29*) (**Fig. 2D & 2E**). We subsequently preformed cryo-immunolabelling using ChAdOx1 nCoV-19 vaccinated mice sera which confirmed the presentation of abundant S protein on the cell surface, but not on control cells infected with ChAdOx-GFP or uninfected cells (**Sup. Fig. 3**).

**Figure 2.**
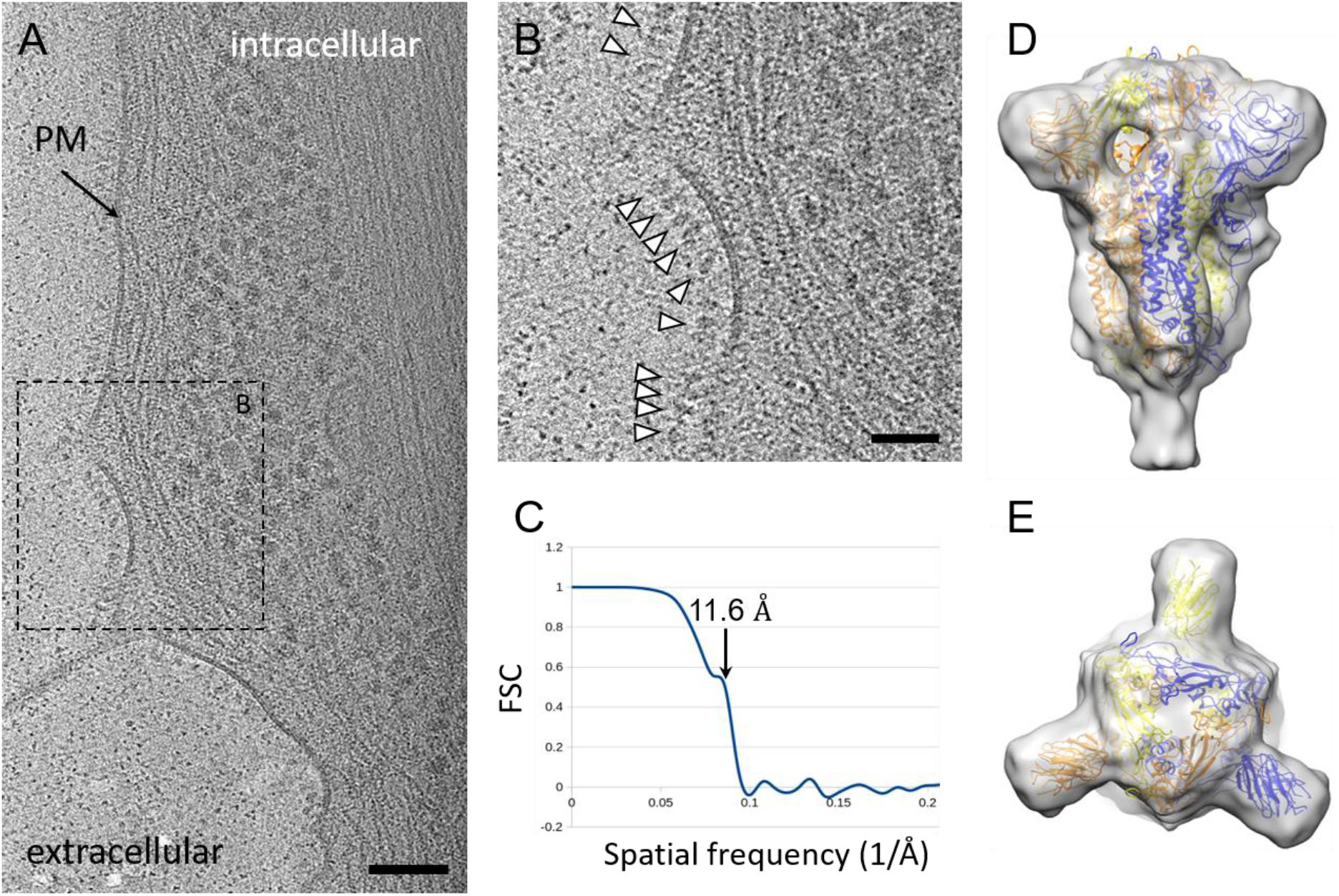
Cryo-ET and subtomogram average of ChAdOx1 nCoV-19 derived spike. **(A)** Tomographic slice of U2OS cell transduced with ChAdOx1 nCoV-19. The slice is 6.4 Å thick; PM = plasma membrane, scale bar = 100 nm **(B)** Detailed view of the boxed area marked in **(A)**. White arrowheads indicate spike proteins on the cell surface; scale bar = 50 nm. **(C-E)** Subtomogram average of ChAdOx1 nCoV-19 spikes at 11.6 Å resolution as indicated by Fourier-Shell correlation at 0.5 cut-off **(C)**, shown from side view **(D)**, and top view **(E).** SARS-CoV-2 atomic model (PDB 6ZB5) (*29*) is fitted for reference.

**Figure 3:**
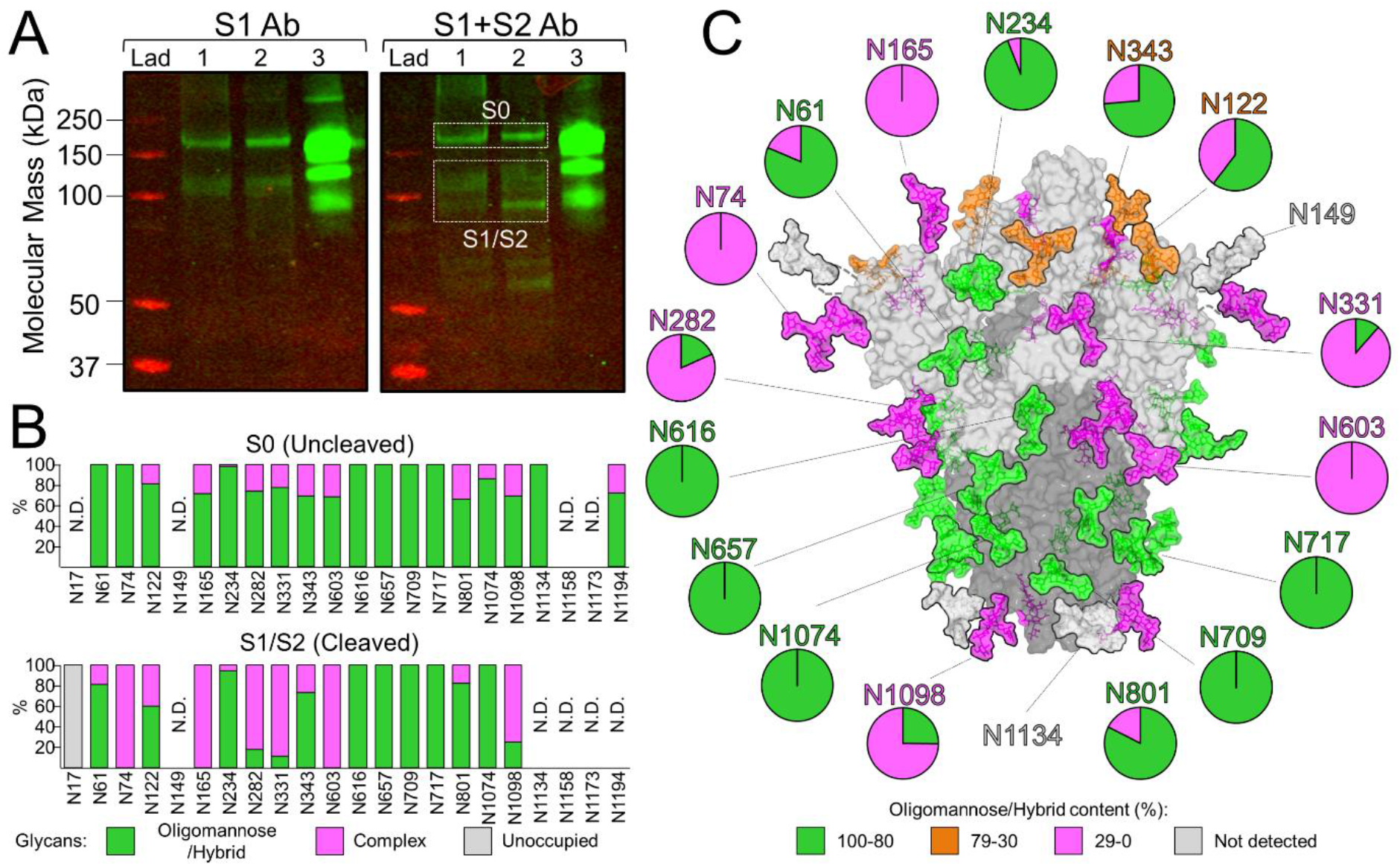
Site-specific glycan processing of SARS-CoV-2 S upon infection with ChAdOx1 nCoV-19. **(A)** Western blot analysis of SARS-CoV-2 spike proteins, using anti-S1 and anti-S1+S2 antibodies. Lane 1= Protein pellet from 293F cell lysates infected with ChAdOx1 nCoV-19. Lane 2= Reduced protein pellet from 293F infected with ChAdOx1 nCoV-19. Lane 3=2P-stablilsed SARS-CoV-2 S protein. The white boxes correspond to gel bands that were excised for mass spectrometric analysis. **(B)** Site-specific N-linked glycosylation of SARS-CoV-2 S0 and S1/S2 glycoproteins. The bar graphs represent the relative quantities of digested glycopeptides possessing the identifiers of oligomannose/hybrid-type glycans (green), complex-type glycans (pink), unoccupied PNGs (grey), or not determined (N.D.) at each N-linked glycan sequon on the S protein, listed from N to C terminus. **(C)** Glycosylated model of the cleaved (S1/S2) SARS-CoV-2 spike. The pie charts summarise the mass spectrometric analysis of the oligomannose/hybrid (green), complex (pink), or unoccupied (grey) N-linked glycan populations. Representative glycans are modelled onto the prefusion structure of trimeric SARS-CoV-2 S glycoprotein (PDB ID: 6VSB) (*3*), with one RBD in the “up” conformation. The modelled glycans are coloured according to oligomannose/hybrid-glycan content with glycan sites labelled in green (80-100%), orange (30-79%), pink (0-29%) or grey (not detected).

In order to investigate the presence of post-fusion S protein conformations, we employed template searching for pre- and post-fusion spikes on cells infected with ChAdOx nCoV-19 (**Sup. Fig. 4**). This analysis revealed a presence of abundant pre-fusion spikes on the cell surface, while little post-fusion spikes are detected. Given the lack of a post-fusion conformation specific anti-S2 antibody, we were not able to probe, by cryo-immunolabelling, post-fusion spikes on the surface of cells.

The presentation of prefusion S proteins was achieved through encoding SARS-CoV-2 S protein in the ChAdOx1 backbone without the incorporation of stabilising mutations. Given that most neutralizing antibodies target epitopes displayed on the prefusion spike, the cryoET analysis revealing the expression of trimeric SARS-CoV-2 spike in the prefusion conformation strongly supports ChAdOx nCoV-19 as an effective vaccine strategy for the generation of a neutralizing immune response.

### Site-specific glycan analysis of the ChAdOx1 nCoV-19 derived SARS-CoV-2 S protein reveals native-like glycan maturation

The SARS-CoV-2 S gene encodes 22 N-linked glycosylation sequons which span both the S1 and S2 subunits (**Fig. 1A**). These host-derived glycans mask immunogenic protein epitopes from the humoral immune system; a common strategy utilised by viruses to evade the immune system (*30*). It is therefore of considerable importance that spike proteins produced upon vaccination successfully recapitulates the glycosylation observed on the virus, since antibodies may be elicited against protein epitopes that are occluded during natural infection. We sought to resolve the site-specific glycosylation of SARS-CoV-2 S proteins produced by cells following infection with ChAdOx1 nCoV-19. To this end, human embryonic kidney 293 F (HEK293F) cells were incubated with ChAdOx1 nCoV-19. Western blot analysis of the protein pellets from the cell lysates using both S1- and S2-specific antibodies confirmed the presence and maturation of the S protein into the S1 and S2 subunits following furin cleavage. Since the whole cell was lysed, intracellular material of uncleaved S0 protein was also detected (**Fig. 3A**). These gel bands were excised and analysed by liquid-chromatography-mass spectrometry (LC-MS) separately.

To determine the site-specific glycosylation and glycan occupancy of the spike protein, we employed trypsin, chymotrypsin, alpha-lytic protease to generate glycopeptide samples. In order to increase the signal intensity of the glycopeptides, these samples were treated sequentially with Endoglycosidase H to cleave oligomannose- and hybrid-type glycans between the core GlcNAc residues, leaving a single GlcNAc attached to the Asn residue (+203Da). Subsequent PNGase F treatment removed the remaining complex-type glycans which deamidates the Asn to Asp resulting in a +3 Da shift as the reaction was conducted in O^18^ water (*31*–*33*). This homogenisation of the glycans into oligomannose/hybrid-type and complex-type glycan categories was performed in order to increase coverage of across the S protein. The mass of peptides with unoccupied glycan sites exhibit no mass shift. The resulting peptide/glycopeptide pool from both the S1/S2 (cleaved) and S0 (uncleaved) proteins were subjected to LC-MS analysis (**Fig. 3B**).

The glycans presented on this S0 immature form of the protein were predominantly underprocessed oligomannose/hybrid-type glycans (85%) (**Sup. Table 1**), which reflects the intracellular origin of the protein, yet to be trafficked through to the cell membrane. This is consistent with furin protease being located in the *trans*-Golgi apparatus. In contrast, the cleaved S1/S2 protein, likely to be presented on the surface of cells, possesses significantly lower levels of these underprocessed glycans (56%) and elevated levels of complex-type glycans (38%). Furthermore, high levels of glycan occupancy were observed across the protein. This is especially important since glycans are known to shield more immunogenic protein epitopes, thus the expression of SARS-CoV-2 S proteins using the ChAdOx1 vaccine platform results in presentation of epitopes similar to those seen during natural infection (*5*, *30*, *34*). We note that the overall levels of oligomannose/hybrid-type glycans on the S1/S2 protein were elevated compared previous analyses, however we attribute this to the lack of coverage of the glycosylation sites towards the C-terminus which are predominantly complex-type on recombinant proteins and viruses (*5*, *27*, *35*).

Using the cryo-EM structure of the trimeric SARS-CoV-2 S protein, we mapped the glycosylation status of the S1/S2 protein (**Fig. 3C**). A mixture of oligomannose/hybrid and complex-type sites were observed, with glycan sites such as N234, which are known to have stabilising effects on the RBD, preserving the predominantly oligomannose state reported in both recombinant proteins and viruses (*5*, *27*, *35*). Similarly, the glycan at N165 which also stabilises the RBD “up” conformation was determined to be complex-type on the S protein arising from infection of cells with ChAdOx1 nCoV19. Since glycans are sensitive reporters of local protein architecture, it is encouraging that such glycans, known to have structural roles, conserve their processing state which provides additional evidence of native-like prefusion protein structure.

Adenoviruses have broad cell tropism due to the widespread distribution of their cellular receptors such as coxsackie and adenovirus receptor (CAR) and CD46 (*36*, *37*). Replication-deficient adenovirus vectored vaccines, such as ChAdOx nCoV-19, are administered intramuscularly and predominantly induce antigen expression in muscle cells, fibroblasts and professional antigen presenting cells (APCs, including dendritic cells) present in germinal centres of the lymph nodes. These APCs are both primed by direct adenovirus infection or by cross-priming from non-immune cells (*37*).

During vaccination, SARS-CoV-2 S protein processing can be dependent on both the receptors present on, and the enzymes expressed by the S-producing cells (*38*). Presence of the cell entry receptor ACE2 and host-cell proteases including furin and TMPRSS2 can drive conformational rearrangements resulting in post-fusion conformations of S. Tissue distribution of ACE2 is primarily in the nasal mucosa and GI tracts, and TMPRSS2 is predominantly expressed in lungs and GI tracts, where cells are susceptible to SARS-CoV-2 infection. In contrast, the muscle, fibroblast and immune cell types targeted by intramuscularly administered ChAdOx1 nCoV-19, have not been shown to express high levels of ACE2 or TMPRSS2 (*39*–*41*). The cell lines chosen for this study of the higher order structure and conformation vaccine-derived S (HeLa S3 and U20S) similarly do not express these enzymes in high abundance and are therefore a suitable surrogate for this *in vitro* study. Although HEK293F cells are not a natural host cell type in which the S protein would be expressed upon administration of ChAdOx nCoV-19, due to the conservation of the enzymatic processing of the early stages of the mammalian N-linked glycosylation pathway (*42*, *43*), they represent a suitable model system to study the nature of the carbohydrate modifications of the S protein. It may be useful in future to examine vaccine derived antigen processing in primary cells or even biopsy material, especially if alternative routes, such as intra-nasal administration, are considered.

Overall, this study provides holistic biophysical scrutiny of the spike protein that is produced upon vaccination with ChAdOx1 nCoV-19. Importantly, we reveal the native-like mimicry of SARS-CoV-2 S protein’s receptor binding functionality, prefusion structure, and processing of glycan modifications. Whilst there is some evidence of S1 subunit shedding, as observed on the actual SARS-CoV-2 virus, the impact of the catabolism of the trimeric spike on immunogenicity and vaccine efficacy is yet to be determined. The data presented here will assist comparison across SARS-CoV-2 vaccination strategies and aid the development of next-generation immunogens.

## Materials and Methods

### Production of ChAdOx1 nCoV-19

ChAdOx1 nCoV-19 was produced as described previously (*17*). The spike protein (S) of SARS-Cov-2 (Genbank accession number YP_009724390.1) was codon optimised for expression in human cell lines and synthesised by GeneArt Gene Synthesis (Thermo Fisher Scientific). The sequence encoding amino acids 2-1273 were cloned into a shuttle plasmid following InFusion cloning (Clontech). The shuttle plasmid encodes a modified human cytomegalovirus major immediate early promoter (IE CMV) with tetracycline operator (TetO) sites, poly adenylation signal from bovine growth hormone (BGH) and a tPA signal sequence upstream of the inserted gene.

### Production of ChAdOx1 nCoV-19 derived SARS-CoV-2 S

A litre of human embryonic kidney 293 Freestyle (HEK293F) cells, at 1×10^6^ cells/ml density, were infected with ChAdOx1 nCoV-19 at an MOI of ~1 viral particles per cell. The cells were incubated at 37 °C for 48 hours and pelleted in a centrifuge. The pellets were washed twice with phosphate buffered saline before addition of 5 ml lysis buffer (20 mM Tris-HCl (pH 8), 150 mM NaCl 0.5% IGEPAL, 0.25% sodium deoxycholate, 0.1% sodium dodecyl sulfate, 1mM EDTA) and sonication. The mixture was centrifuged at 4,000 rpm for 30 mins and the supernatant was collected. 5 ml of lysis buffer and 3.5 ml of chloroform were added, and the tube was vortexed. 9 ml H2O was added and after vortexing, the tube was centrifuged at 4,000 rpm for 30 mins. The upper liquid layer was removed, and 9 ml of methanol was added and after vortexing, the tube was centrifuged again. The methanol was subsequently removed, and the pellet was allowed to air dry in the dark. Pellets were dissolved in 50 mM sodium phosphate, 300 mM sodium chloride, pH 7.

### Expression and purification of recombinant 2P-stabilised SARS-CoV-2 S

Recombinant 2P-stabilised SARS-CoV-2 S protein was expressed as previously described (*3*, *5*). In brief, the expression vector containing the 2P-stabilised S protein was used to transiently transfect FreeStyle293F cells. The protein was purified from the supernatant using nickel-affinity chromatography before size-exclusion chromatography.

### Western blot analysis

Protein pellets from the HEK293F cell lysates were loaded onto an SDS-PAGE gel, then transferred to a PVDF membrane. Polyclonal rabbit anti-S1 antibody (Sinobiological, Cat: 40592-T62) and anti-S2 antibody (Abcam, Cat: ab272504) were used as primary antibodies.

### FACS analysis for cell surface spike expression

Hela S3 cells at 4 ×10^5^ cell/ml were infected with 10 MOI ChAdOx1 nCoV-19 and incubated at 37°C for 40-48h. Cells were harvested and pelleted by centrifugation at 500 g for 5 min. Resuspended cells were split into three pools for the alternative staining and were incubated with either pooled prime-boost sera derived from outbred CD1 mice immunised intramuscularly with 10^8^ infectious units of ChAdOx1 nCoV-19 (*18*), prepared at 1:50 in PBS-BSA-0.5% or human mAbs prepared at 1 μg/ml or recombinant ACE2 expressed with a human Fc tag prepared at 2 μg/ml (see below). Cells were incubated for 2h at RT. Cells were washed 3 × with PBS-BSA-0.5% then incubated for 1h at RT with secondary antibodies conjugated with Alexafluor488 diluted at 1:1000 in PBS-BSA 0.5% (for ChAdOx1 nCov-19 serum staining Goat Anti mouse AlexaFluor 488 (Life Tech A11029), for ACE2 staining, Goat Anti Human AlexaFluor 488 (Life Tech)). Cells were washed 2 × with PBS-BSA-0.5% before resuspending in PBS and analysing by flow cytometry using a Fortessa X20 FACS analyser. Samples were considered positive for spike expression if they had a fluorescence intensity above a threshold value determined by the maximum intensity of the non-infected control cells (Fig 1.D). Experiments were performed twice in duplicate, representative data is shown. Data were analysed using FlowJo v9 (TreeStar).

### ACE2 expression

The ectodomain of the human angiotensin-converting enzyme 2 (ACE2) soluble construct encoding residues 18-740 (from NCBI Reference Sequence: NP_001358344.1.), where the transmembrane and cytoplasmic domains were removed, was expressed using human embryonic kidney 293 F cells (HEK293F). A N-terminal monoFc followed by TEV cleavage was included as well as a C-terminal four amino acid C-tag (EPEA) for affinity purification. ACE2 ectodomain was transiently expressed in Expi293™ (Thermo Fisher Scientific) and protein purified from culture supernatants by C-tag affinity purification followed by gel filtration in Tris-buffered saline (TBS) pH 7.4 buffer.

### Isolation of human monoclonal antibodies from peripheral B cells by spike-specific single B cells sorting

To isolate Spike-specific B cells, PBMCs were labelled with recombinant trimeric spike-twin-Strep and stained with antibody cocktail consisting of CD3-FITC, CD14-FITC, CD56-FITC, CD16-FITC, IgM-FITC, IgA-FITC, IgD-FITC, IgG-BV786, CD19-BUV395 and Strep-MAB-DY549 (iba) to probe the Strep tag of spike. CD19+, IgG+, CD3-, CD14-, CD56-, CD16-,IgM-, IgA-, IgD-, Spike+ cells were then single cell sorted into 96-well PCR plates containing RNase inhibitor (N2611; Promega) and stored at −80 °C.

Genes encoding Ig VH, Ig Vκ and Vλ from positive wells were amplified by RT-PCR (210210; QIAGEN) and nested PCR (203205; Qiagen) using ‘cocktails’ of primers specific for human IgG. PCR products of genes encoding heavy and light chains were joined with the expression vector for human IgG1 or Ig κ-chain or λ-chain (gifts from H. Wardemann) by Gibson assembly. Plasmids encoding heavy and light chains were then co-transfected into the 293T cell line by the polyethylenimine method (408727; Sigma), and antibody were harvested for further study.

### CryoET sample preparation and imaging

EM grids (G300F1, R2/2 Quantifoil holey carbon, gold) were glow-discharged and treated with bovine fibronectin (20 μg/ml) for 30 minutes. Grids were washed with PBS and UV-treated for 1h. 1.6×10^5^ of U2OS or HeLa cells resuspended in 2 ml of DMEM 10% FBS, Pen/Strep were seeded on top of the grids in, in 6 well plate wells. Cells and grids were incubated for 24h at 37°C /5% CO2 to allow cell attachment to grid carbon. ChAdOx1 nCoV-19 was added to the cells at MOI 1 and cells were returned to 37°C/5% CO2 for 48h. Grids were plunge-frozen in liquid ethane at −183°C using the Leica GP2 plunger, after receiving 1 μl of 5× concentrated 10 nm Au fiducial solution (EMS) in the back-side and being blotted from the back. Grids were stored at liquid nitrogen until time of imaging.

Tilt series acquisition was carried out at a FEI Titan Krios G2 (Thermo Fisher Scientific) electron microscope operated at 300 kV and equipped with a Gatan BioQuantum energy filter and post-GIF K3 detector (Gatan, Pleasanton, CA) housed at electron Bioimaging Center (eBIC/Diamond Light Source, UK). Tilt series were acquired using SerialEM (*44*) at a pixel size of 2.13 Å. Zero-loss imaging was used for all tilt series with a 20 eV slit width. Defocus values ranged from −2 μm to −7 μm.

A grouped dose-symmetric scheme was used for all tilt-series (*45*); with a range of +/−60 degrees at 3 degree increments in groups of 3 and total dose of 120-135 e^-^/Å2. At each tilt, 5 movie frames were recorded using Correlated Double Sampling (CDS) in super-resolution mode and saved in lzw compressed tif format with no gain normalisation. Movies were subsequently gain normalised during motion correction and Fourier cropped back to physical pixel size with MotionCor2 (*46*). Tilt-series were manually aligned using eTomo (*47*) and tilt-series and alignment files were imported to emClarity (*25*). 11391 subtomograms were selected after template matching using EMDB-21452 (*4*) as template, low-pass filtered to 40 Å. Subtomograms were separated in 2 completely independent half-datasets and iteratively aligned and averaged resulting in a final 11.6 Å map at 0.5 FSC threshold. Model fitting and visualization were performed in Chimera (*48*).

### Glycopeptide analysis by liquid chromatography-mass spectrometry

Approximately 90 gel bands containing the furin cleaved (S1/S2) and uncleaved (S0) SARS-CoV-2 S protein were excised in order to gather sufficient material. The gel bands were de-stained in 50% 100 mM ammonium bicarbonate, 50% acetonitrile overnight. In-gel reduction and alkylation were performed according to the protocol by Shevchenko *et al.* (*49*). The proteins were digested using trypsin, chymotrypsin and alpha-lytic protease, which cleave at the amino acids R/K, F/Y/W, and T/A/S/V, respectively. After extraction, the peptide/glycopeptides were first treated with endoglycosidase H to deplete oligomannose-type glycans and leave a single GlcNAc residue at the corresponding site. Subsequently, the reaction mixture was completely dried and resuspended in a mixture containing PNGase F using only H2O^18^ (Sigma-Aldrich) throughout. This reaction cleaves the remaining complex-type glycans but leaves the GlcNAc residues remaining after Endo H digestion intact. The use of H2O^18^ enables complex-type glycan sites to be differentiated from unoccupied glycan sites since the hydrolysis of the glycosidic bond by PNGase F leaves an O^18^ isotope on the resulting aspartic acid residue. The resulting peptides are purified using C18 Zip-tip (MerckMillipore) clean-up following the manufacturer’s protocol. Eluted glycopeptides were dried and re-suspended in 0.1% formic acid prior to analysis by liquid chromatography-mass spectrometry. An Easy-nLC 1200 (Thermo Fisher Scientific) system coupled to an Orbitrap Fusion mass spectrometer (Thermo Fisher Scientific) using higher energy collision-induced dissociation fragmentation was used. Peptides were separated using an EasySpray PepMap RSLC C18 column (75 cm × 75 μm) with a 275-minute linear gradient consisting of 0-32% acetonitrile in 0.1% formic acid over 240 minutes, followed by 35 minutes of 80% acetonitrile in 0.1% formic acid. The flow rate was set to 200 nl/min. The spray voltage was set to 2.7 kV and the temperature of the heated capillary was set to 40 °C. The ion transfer tube temperature was set to 275 °C. The scan range was 400-1600 m/z. The HCD collision energy was set to 27%. Precursor and fragment detection were performed at a resolution of MS^1^=100,000 and MS^2^=30,000. The AGC target for MS^1^=4e^5^ and MS^2^=5e^4^ and injection time for MS^1^= 50 ms and MS^2^= 54 ms.

Glycopeptide fragmentation data were extracted from the raw file using Byonic™ (Version 3.5) and Byologic™ software (Version 3.5; Protein Metrics Inc.). The glycopeptide fragmentation data were evaluated manually for each glycopeptide; the peptide was scored as true-positive when the correct b- and y-fragment ions were observed. Two modifications were searched for: +203 Da corresponding to a single GlcNAc residue, still attached to an asparagine residue of an N-linked glycan site, following the cleavage by Endo H, and +3 D corresponding to the O^18^ deamidation product of a complex glycan. The relative quantities of each glycan type at each site as well as the unoccupied proportion were determined by comparison of the extracted ion chromatographic areas.

### Glycosylated model construction

Structural models of N-linked glycan presentation on SARS-CoV-2 S were created using electron microscopy structures (PDB ID:6VSB) along with complex-, hybrid-, an oligomannose-type N-linked glycans (PDB ID 4BYH, 4B7I, and 2WAH). A representative glycoform presented at each site was modelled on to the N-linked carbohydrate attachment sites in Coot (*50*).

## Author Contributions

Y.W. T.L, S.G., M. C. and P.Z. conceived the research and designed the experiments. L.M. prepared cell culture, cryoEM samples and with A.K. performed cryoET data processing and subtomogram averaging. A.H. collected cryoET data and performed initial tomography reconstruction. E.R.A. performed FACS experiments and analyzed data. D.P., F.D., H.D. and H.C. cloned, expressed and purified proteins in this work. G.R.S., J.M, W.D, P.S. provided human monoclonal antibodies. Y.W. and J.A. performed mass spectrometry experiments and analyzed data. Y.W., L.M., E.R.A., T.L., S.G., M.C. and P.Z. analyzed data and wrote the paper. All authors commented on and approved the final manuscript.

## Data and materials availability

The cryoEM density maps for ChAdOx nCoV-2019 was deposited in the EMDB under accession code XXX. Mass spectrometry raw files have been deposited in the MassIVE proteomics database (DOI: XXX).

## References

1. F. Amanat, F. Krammer, SARS-CoV-2 Vaccines: Status Report. Immunity. 52 (2020), pp. 583–589.

2. M. Letko, A. Marzi, V. Munster, Functional assessment of cell entry and receptor usage for SARS-CoV-2 and other lineage B betacoronaviruses. Nat. Microbiol., 1–8 (2020).

3. D. Wrapp, N. Wang, K. S. Corbett, J. A. Goldsmith, C.-L. Hsieh, O. Abiona, B. S. Graham, J. S. McLellan, Cryo-EM structure of the 2019-nCoV spike in the prefusion conformation. Science (2020), doi:10.1126/science.abb2507.

4. A. C. Walls, Y. J. Park, M. A. Tortorici, A. Wall, A. T. McGuire, D. Veesler, Structure, Function, and Antigenicity of the SARS-CoV-2 Spike Glycoprotein. Cell. 181, 281–292.e6 (2020).

5. Y. Watanabe, J. D. Allen, D. Wrapp, J. S. McLellan, M. Crispin, Site-specific glycan analysis of the SARS-CoV-2 spike. Science (2020), doi:10.1126/science.abb9983.

6. F. Krammer, SARS-CoV-2 vaccines in development. Nature, 1–16 (2020).

7. F. P. Polack, S. J. Thomas, N. Kitchin, J. Absalon, A. Gurtman, S. Lockhart, J. L. Perez, G. Pérez Marc, E. D. Moreira, C. Zerbini, R. Bailey, K. A. Swanson, S. Roychoudhury, K. Koury, P. Li, W. V. Kalina, D. Cooper, R. W. Frenck, L. L. Hammitt, Ö. Türeci, H. Nell, A. Schaefer, S. Ünal, D. B. Tresnan, S. Mather, P. R. Dormitzer, U. Şahin, K. U. Jansen, W. C. Gruber, Safety and Efficacy of the BNT162b2 mRNA Covid-19 Vaccine. N. Engl. J. Med. 383, 2603–2615 (2020).

8. K. S. Corbett, B. Flynn, K. E. Foulds, J. R. Francica, S. Boyoglu-Barnum, A. P. Werner, B. Flach, S. O’Connell, K. W. Bock, M. Minai, B. M. Nagata, H. Andersen, D. R. Martinez, A. T. Noe, N. Douek, M. M. Donaldson, N. N. Nji, G. S. Alvarado, D. K. Edwards, D. R. Flebbe, E. Lamb, N. A. Doria-Rose, B. C. Lin, M. K. Louder, S. O’Dell, S. D. Schmidt, E. Phung, L. A. Chang, C. Yap, J.-P. M. Todd, L. Pessaint, A. Van Ry, S. Browne, J. Greenhouse, T. Putman-Taylor, A. Strasbaugh, T.-A. Campbell, A. Cook, A. Dodson, K. Steingrebe, W. Shi, Y. Zhang, O. M. Abiona, L. Wang, A. Pegu, E. S. Yang, K. Leung, T. Zhou, I.-T. Teng, A. Widge, I. Gordon, L. Novik, R. A. Gillespie, R. J. Loomis, J. I. Moliva, G. Stewart-Jones, S. Himansu, W.-P. Kong, M. C. Nason, K. M. Morabito, T. J. Ruckwardt, J. E. Ledgerwood, M. R. Gaudinski, P. D. Kwong, J. R. Mascola, A. Carfi, M. G. Lewis, R. S. Baric, A. McDermott, I. N. Moore, N. J. Sullivan, M. Roederer, R. A. Seder, B. S. Graham, Evaluation of the mRNA-1273 Vaccine against SARS-CoV-2 in Nonhuman Primates. N. Engl. J. Med. (2020), doi:10.1056/nejmoa2024671.

9. Q. Gao, L. Bao, H. Mao, L. Wang, K. Xu, M. Yang, Y. Li, L. Zhu, N. Wang, Z. Lv, H. Gao, X. Ge, B. Kan, Y. Hu, J. Liu, F. Cai, D. Jiang, Y. Yin, C. Qin, J. Li, X. Gong, X. Lou, W. Shi, D. Wu, H. Zhang, L. Zhu, W. Deng, Y. Li, J. Lu, C. Li, X. Wang, W. Yin, Y. Zhang, C. Qin, Development of an inactivated vaccine candidate for SARS-CoV-2. Science. 369, 77–81 (2020).

10. C. Liu, L. Mendonça, Y. Yang, Y. Gao, C. Shen, J. Liu, T. Ni, B. Ju, C. Liu, X. Tang, J. Wei, X. Ma, Y. Zhu, W. Liu, S. Xu, Y. Liu, J. Yuan, J. Wu, Z. Liu, Z. Zhang, L. Liu, P. Wang, P. Zhang, The Architecture of Inactivated SARS-CoV-2 with Postfusion Spikes Revealed by Cryo-EM and Cryo-ET. Structure (2020), doi:10.1016/j.str.2020.10.001.

11. S. J. Zost, P. Gilchuk, J. B. Case, E. Binshtein, R. E. Chen, J. P. Nkolola, A. Schäfer, J. X. Reidy, A. Trivette, R. S. Nargi, R. E. Sutton, N. Suryadevara, D. R. Martinez, L. E. Williamson, E. C. Chen, T. Jones, S. Day, L. Myers, A. O. Hassan, N. M. Kafai, E. S. Winkler, J. M. Fox, S. Shrihari, B. K. Mueller, J. Meiler, A. Chandrashekar, N. B. Mercado, J. J. Steinhardt, K. Ren, Y. M. Loo, N. L. Kallewaard, B. T. McCune, S. P. Keeler, M. J. Holtzman, D. H. Barouch, L. E. Gralinski, R. S. Baric, L. B. Thackray, M. S. Diamond, R. H. Carnahan, J. E. Crowe, Potently neutralizing and protective human antibodies against SARS-CoV-2. Nature. 584, 443–449 (2020).

12. D. F. Robbiani, C. Gaebler, F. Muecksch, J. C. C. Lorenzi, Z. Wang, A. Cho, M. Agudelo, C. O. Barnes, A. Gazumyan, S. Finkin, T. Hägglöf, T. Y. Oliveira, C. Viant Hurley, H. H. Hoffmann, K. G. Millard, R. G. Kost, M. Cipolla, K. Gordon, F. Bianchini, S. T. Chen, V. Ramos, R. Patel, J. Dizon, I. Shimeliovich, P. Mendoza, H. Hartweger, L. Nogueira, M. Pack, J. Horowitz, F. Schmidt, Y. Weisblum, E. Michailidis, A. W. Ashbrook, E. Waltari, J. E. Pak, K. E. Huey-Tubman, N. Koranda, P. R. Hoffman, A. P. West, C. M. Rice, T. Hatziioannou, P. J. Bjorkman, P. D. Bieniasz, M. Caskey, M. C. Nussenzweig, Convergent antibody responses to SARS-CoV-2 in convalescent individuals. Nature. 584, 437–442 (2020).

13. S. Du, Y. Cao, Q. Zhu, P. Yu, F. Qi, G. Wang, X. Du, L. Bao, W. Deng, H. Zhu, J. Liu, J. Nie, Y. Zheng, H. Liang, R. Liu, S. Gong, H. Xu, A. Yisimayi, Q. Lv, B. Wang, R. He, Y. Han, W. Zhao, Y. Bai, Y. Qu, X. Gao, C. Ji, Q. Wang, N. Gao, W. Huang, Y. Wang, X. S. Xie, X. dong Su, J. Xiao, C. Qin, Structurally Resolved SARS-CoV-2 Antibody Shows High Efficacy in Severely Infected Hamsters and Provides a Potent Cocktail Pairing Strategy. Cell (2020), doi:10.1016/j.cell.2020.09.035.

14. C. O. Barnes, C. A. Jette, M. E. Abernathy, K.-M. A. Dam, S. R. Esswein, H. B. Gristick, A. G. Malyutin, N. G. Sharaf, K. E. Huey-Tubman, Y. E. Lee, D. F. Robbiani, M. C. Nussenzweig, A. P. West, P. J. Bjorkman, SARS-CoV-2 neutralizing antibody structures inform therapeutic strategies. Nature, 1–9 (2020).

15. P. J. M. Brouwer, T. G. Caniels, K. van der Straten, J. L. Snitselaar, Y. Aldon, S. Bangaru, J. L. Torres, N. M. A. Okba, M. Claireaux, G. Kerster, A. E. H. Bentlage, M. M. van Haaren, D. Guerra, J. A. Burger, E. E. Schermer, K. D. Verheul, N. van der Velde, A. van der Kooi, J. van Schooten, M. J. van Breemen, T. P. L. Bijl, K. Sliepen, A. Aartse, R. Derking, I. Bontjer, N. A. Kootstra, W. J. Wiersinga, G. Vidarsson, B. L. Haagmans, A. B. Ward, G. J. de Bree, R. W. Sanders, M. J. van Gils, Potent neutralizing antibodies from COVID-19 patients define multiple targets of vulnerability. Science, eabc5902 (2020).

16. P. M. Folegatti, K. J. Ewer, P. K. Aley, B. Angus, S. Becker, S. Belij-Rammerstorfer, D. Bellamy, S. Bibi, M. Bittaye, E. A. Clutterbuck, C. Dold, S. N. Faust, A. Finn, A. L. Flaxman, B. Hallis, P. Heath, D. Jenkin, R. Lazarus, R. Makinson, A. M. Minassian, K. M. Pollock, M. Ramasamy, H. Robinson, M. Snape, R. Tarrant, M. Voysey, C. Green, A. D. Douglas, A. V. S. Hill, T. Lambe, S. C. Gilbert, A. J. Pollard, Oxford COVID Vaccine Trial Group, Safety and immunogenicity of the ChAdOx1 nCoV-19 vaccine against SARS-CoV-2: a preliminary report of a phase 1/2, single-blind, randomised controlled trial. Lancet. 0 (2020), doi:10.1016/S0140-6736(20)31604-4.

17. N. van Doremalen, T. Lambe, A. Spencer, S. Belij-Rammerstorfer, J. N. Purushotham, J. R. Port, V. A. Avanzato, T. Bushmaker, A. Flaxman, M. Ulaszewska, F. Feldmann, E. R. Allen, H. Sharpe, J. Schulz, M. Holbrook, A. Okumura, K. Meade-White, L. Pérez-Pérez, N. J. Edwards, D. Wright, C. Bissett, C. Gilbride, B. N. Williamson, R. Rosenke, D. Long, A. Ishwarbhai, R. Kailath, L. Rose, S. Morris, C. Powers, J. Lovaglio, P. W. Hanley, D. Scott, G. Saturday, E. de Wit, S. C. Gilbert, V. J. Munster, ChAdOx1 nCoV-19 vaccine prevents SARS-CoV-2 pneumonia in rhesus macaques. Nature, 1–5 (2020).

18. S. P. Graham, R. K. McLean, A. J. Spencer, S. Belij-Rammerstorfer, D. Wright, M. Ulaszewska, J. C. Edwards, J. W. P. Hayes, V. Martini, N. Thakur, C. Conceicao, I. Dietrich, H. Shelton, R. Waters, A. Ludi, G. Wilsden, C. Browning, D. Bialy, S. Bhat, P. Stevenson-Leggett, P. Hollinghurst, C. Gilbride, D. Pulido, K. Moffat, H. Sharpe, E. Allen, V. Mioulet, C. Chiu, J. Newman, A. S. Asfor, A. Burman, S. Crossley, J. Huo, R. J. Owens, M. Carroll, J. A. Hammond, E. Tchilian, D. Bailey, B. Charleston, S. C. Gilbert, T. J. Tuthill, T. Lambe, Evaluation of the immunogenicity of prime-boost vaccination with the replication-deficient viral vectored COVID-19 vaccine candidate ChAdOx1 nCoV-19. npj Vaccines. 5, 1–6 (2020).

19. M. N. Ramasamy, A. M. Minassian, K. J. Ewer, A. L. Flaxman, P. M. Folegatti, D. R. Owens, M. Voysey, P. K. Aley, B. Angus, G. Babbage, S. Belij-Rammerstorfer, L. Berry, S. Bibi, M. Bittaye, K. Cathie, H. Chappell, S. Charlton, P. Cicconi, E. A. Clutterbuck, R. Colin-Jones, C. Dold, K. R. W. Emary, S. Fedosyuk, M. Fuskova, D. Gbesemete, C. Green, B. Hallis, M. M. Hou, D. Jenkin, C. C. D. Joe, E. J. Kelly, S. Kerridge, A. M. Lawrie, A. Lelliott, M. N. Lwin, R. Makinson, N. G. Marchevsky, Y. Mujadidi, A. P. S. Munro, M. Pacurar, E. Plested, J. Rand, T. Rawlinson, S. Rhead, H. Robinson, A. J. Ritchie, A. L. Ross-Russell, S. Saich, N. Singh, C. C. Smith, M. D. Snape, R. Song, R. Tarrant, Y. Themistocleous, K. M. Thomas, T. L. Villafana, S. C. Warren, M. E. E. Watson, A. D. Douglas, A. V. S. Hill, T. Lambe, S. C. Gilbert, S. N. Faust, A. J. Pollard, J. Aboagye, K. Adams, A. Ali, E. Allen, L. Allen, J. Allison, F. Andritsou, R. Anslow, E. H. Arbe-Barnes, M. Baker, N. Baker, P. Baker, I. Baleanu, D. Barker, E. Barnes, J. R. Barrett, K. Barrett, L. Bates, A. Batten, K. Beadon, R. Beckley, D. Bellamy, A. Berg, L. Bermejo, E. Berrie, A. Beveridge, K. R. Bewley, E. M. Bijker, G. Birch, L. Blackwell, H. Bletchly, C. Blundell, S. Blundell, E. Bolam, E. Boland, D. Bormans, N. Borthwick, K. Boukas, T. Bower, F. Bowring, A. Boyd, T. Brenner, P. Brown, C. Brown-O’Sullivan, S. Bruce, E. Brunt, J. Burbage, J. Burgoyne, K. R. Buttigieg, N. Byard, I. C. Puig, S. Camara, M. Cao, F. Cappuccini, M. Carr, M. W. Carroll, P. Cashen, A. Cavey, J. Chadwick, R. Challis, D. Charles, J. Cho, L. Cifuentes, E. Clark, S. Collins, C. P. Conlon, N. S. Coombes, R. Cooper, C. Cooper, W. E. M. Crocker, S. Crosbie, D. Cullen, C. J. Cunningham, F. Cuthbertson, B. E. Damratoski, L. Dando, M. S. Datoo, C. Datta, H. Davies, S. Davies, E. Davis, J. Davis, D. Dearlove, T. Demissie, S. Di Marco, C. Di Maso, D. DiTirro, C. Docksey, T. Dong, F. R. Donnellan, N. Douglas, C. Downing, J. Drake, R. Drake-Brockman, R. E. Drury, S. J. Dunachie, C. J. Edwards, N. J. Edwards, O. El Muhanna, S. C. Elias, R. S. Elliott, M. J. Elmore, M. R. English, S. Felle, S. Feng, C. Ferreira Da Silva, S. Field, R. Fisher, C. Fixmer, K. J. Ford, J. Fowler, E. Francis, J. Frater, J. Furze, P. Galian-Rubio, C. Galloway, H. Garlant, M. Gavrila, F. Gibbons, K. Gibbons, C. Gilbride, H. Gill, K. Godwin, K. Gordon-Quayle, G. Gorini, L. Goulston, C. Grabau, L. Gracie, N. Graham, N. Greenwood, O. Griffiths, G. Gupta, E. Hamilton, B. Hanumanthadu, S. A. Harris, T. Harris, D. Harrison, T. C. Hart, B. Hartnell, L. Haskell, S. Hawkins, J. A. Henry, M. H. Herrera, D. Hill, J. Hill, G. Hodges, S. H. C. Hodgson, K. Horton, E. Howe, N. Howell, J. Howes, B. Huang, J. Humphreys, H. Humphries, P. Iveson, F. Jackson, S. Jackson, S. Jauregui, H. Jeffers, B. Jones, C. E. Jones, E. Jones, K. Jones, A. Joshi, R. Kailath, J. Keen, E. J. Kelly, D. M. Kelly, S. Kelly, D. Kelly, D. Kerr, L. Khan, A. Killen, J. Kinch, L. D. W. King, T. B. King, L. Kingham, P. Klenerman, J. C. Knight, D. Knott, S. Koleva, G. Lang, C. W. Larkworthy, J. P. J. Larwood, R. L. Law, A. Lee, K. Y. N. Lee, E. A. Lees, S. Leung, Y. Li, A. M. Lias, A. Linder, S. Lipworth, S. Liu, X. Liu, S. Lloyd, L. Loew, R. Lopez Ramon, M. Madhavan, D. Mainwaring, G. Mallett, K. Mansatta, S. Marinou, P. Marius, E. Marlow, P. Marriott, J. L. Marshall, J. Martin, S. Masters, J. McEwan, J. McGlashan, L. McInroy, N. McRobert, C. Megson, A. J. Mentzer, N. Mirtorabi, C. Mitton, M. Moore, M. Moran, E. Morey, R. Morgans, S. R. Morris, H. Morrison, G. Morshead, R. Morter, J. Muller, C. Munro, S. Murphy, P. Mweu, A. Noé, F. L. Nugent, K. O’Brien, D. O’Connor, B. Oguti, V. Olchawski, C. Oliveria, P. J. O’Reilly, P. Osborne, L. Owen, N. Owino, P. Papageorgiou, H. Parracho, H. Parsons, B. Patel, M. Patrick-Smith, Y. Peng, E. Penn, M. P. Peralta Alvarez, J. Perring, C. Petropoulos, D. J. Phillips, D. Pipini, S. Pollard, I. Poulton, D. Pratt, L. Presland, P. Proud, S. Provstgaard-Morys, S. Pueschel, D. Pulido, R. Rabara, K. Radia, D. Rajapaksa, F. Ramos Lopez, H. Ratcliffe, S. Rayhan, B. Rees, E. Pabon, H. Roberts, I. Robertson, S. Roche, C. S. Rollier, R. Romani, Z. Rose, I. Rudiansyah, S. Sabheha, S. Salvador, H. Sanders, K. Sanders, C. Sayce, A. B. Schmid, E. Schofield, G. Screaton, C. Sedik, R. R. Segireddy, B. Selby, I. Shaik, H. R. Sharpe, A. Shea, S. Silk, L. Silva-Reyes, D. T. Skelly, D. J. Smith, D. C. Smith, N. Smith, A. J. Spencer, L. Spoors, E. Stafford, I. Stamford, L. Stockdale, D. Stockley, L. Stockwell, M. Stokes, L. Strickland, S. Sulaiman, E. Summerton, Z. Swash, A. Szigeti, A. Alaoui, R. Tanner, I. J. Taylor, K. Taylor, U. Taylor, R. te Water Naude, A. Themistocleous, M. Thomas, T. Thomas, A. Thompson, K. Thompson, V. Thornton-Jones, L. Tinh, S. Tonks, J. Towner, N. Tran, J. A. Tree, A. Truby, C. Turner, R. Turner, M. Ulaszewska, R. Varughese, D. Verbart, M. K. Verheul, I. Vichos, L. Walker, M. E. Wand, B. Watkins, J. Welch, A. West, C. White, R. White, P. Williams, M. Woodyer, A. T. Worth, D. Wright, T. Wrin, X. L. Yao, D. A. Zbarcea, D. Zizi, Safety and immunogenicity of ChAdOx1 nCoV-19 vaccine administered in a prime-boost regimen in young and old adults (COV002): a single-blind, randomised, controlled, phase 2/3 trial. Lancet. 0 (2020), doi:10.1016/s0140-6736(20)32466-1.

20. S. Sebastian, A. Flaxman, K. M. Cha, M. Ulaszewska, C. Gilbride, H. Sharpe, E. Wright, A. J. Spencer, S. Dowall, R. Hewson, S. Gilbert, T. Lambe, A multi-filovirus vaccine candidate: Co-expression of Ebola, Sudan, and Marburg antigens in a single vector. Vaccines. 8 (2020), doi:10.3390/vaccines8020241.

21. L. Zhang, C. B. Jackson, H. Mou, A. Ojha, H. Peng, B. D. Quinlan, E. S. Rangarajan Pan, A. Vanderheiden, M. S. Suthar, W. Li, T. Izard, C. Rader, M. Farzan, H. Choe, SARS-CoV-2 spike-protein D614G mutation increases virion spike density and infectivity. Nat. Commun. 11, 1–9 (2020).

22. J. Pallesen, N. Wang, K. S. Corbett, D. Wrapp, R. N. Kirchdoerfer, H. L. Turner, C. A. Cottrell, M. M. Becker, L. Wang, W. Shi, W.-P. Kong, E. L. Andres, A. N. Kettenbach, M. R. Denison, J. D. Chappell, B. S. Graham, A. B. Ward, J. S. McLellan, Immunogenicity and structures of a rationally designed prefusion MERS-CoV spike antigen. Proc. Natl. Acad. Sci. U. S. A. 114, E7348–E7357 (2017).

23. A. C. Walls, M. A. Tortorici, J. Snijder, X. Xiong, B. J. Bosch, F. A. Rey, D. Veesler, Tectonic conformational changes of a coronavirus spike glycoprotein promote membrane fusion. Proc. Natl. Acad. Sci. U. S. A. 114, 11157–11162 (2017).

24. I. Berger, C. Schaffitzel, The SARS-CoV-2 spike protein: balancing stability and infectivity. Cell Res. 30 (2020), pp. 1059–1060.

25. B. A. Himes, P. Zhang, emClarity: software for high-resolution cryo-electron tomography and subtomogram averaging. Nat Methods. 15, 955–961 (2018).

26. B. Turoňová, M. Sikora, C. Schürmann, W. Hagen, S. Welsch, F. Blanc, S. von Bülow, M. Gecht, K. Bagola, C. Hörner, G. van Zandbergen, S. Mosalaganti, A. Schwarz, R. Covino, M. Mühlebach, G. Hummer, J. K. Locker, M. Beck, bioRxiv, in press, doi:10.1101/2020.06.26.173476.

27. H. Yao, Y. Song, Y. Chen, N. Wu, J. Xu, C. Sun, J. Zhang, T. Weng, Z. Zhang, Z. Wu, L. Cheng, D. Shi, X. Lu, J. Lei, M. Crispin, Y. Shi, L. Li, S. Li, Molecular architecture of the SARS-CoV-2 virus. Cell (2020), doi:10.1016/j.cell.2020.09.018.

28. Z. Ke, J. Oton, K. Qu, M. Cortese, V. Zila, L. McKeane, T. Nakane, J. Zivanov, C. J. Neufeldt, B. Cerikan, J. M. Lu, J. Peukes, X. Xiong, H. G. Kräusslich, S. H. W. Scheres, R. Bartenschlager, J. A. G. Briggs, Structures and distributions of SARS-CoV-2 spike proteins on intact virions. Nature, 1–5 (2020).

29. C. Toelzer, K. Gupta, S. K. N. Yadav, U. Borucu, A. D. Davidson, M. Kavanagh Williamson, D. K. Shoemark, F. Garzoni, O. Staufer, R. Milligan, J. Capin, A. J. Mulholland, J. Spatz, D. Fitzgerald, I. Berger, C. Schaffitzel, Free fatty acid binding pocket in the locked structure of SARS-CoV-2 spike protein. Science. 370, 725–730 (2020).

30. Y. Watanabe, T. A. Bowden, I. A. Wilson, M. Crispin, Exploitation of glycosylation in enveloped virus pathobiology. Biochim. Biophys. Acta. 1863, 1480–1497 (2019).

31. L. Cao, M. Pauthner, R. Andrabi, K. Rantalainen, Z. Berndsen, J. K. Diedrich, S. Menis, D. Sok, R. Bastidas, S.-K. R. Park, C. M. Delahunty, L. He, J. Guenaga, R. T. Wyatt, W. R. Schief, A. B. Ward, J. R. Yates, D. R. Burton, J. C. Paulson, Differential processing of HIV envelope glycans on the virus and soluble recombinant trimer. Nat. Commun. 9, 3693 (2018).

32. L. Cao, J. K. Diedrich, D. W. Kulp, M. Pauthner, L. He, S.-K. R. Park, D. Sok, C. Y. Su, C. M. Delahunty, S. Menis, R. Andrabi, J. Guenaga, E. Georgeson, M. Kubitz, Y. Adachi, D. R. Burton, W. R. Schief, J. R. Yates III, J. C. Paulson, Global site-specific N-glycosylation analysis of HIV envelope glycoprotein. Nat. Commun. 8, 14954 (2017).

33. W. B. Struwe, E. Chertova, J. D. Allen, G. E. Seabright, Y. Watanabe, D. J. Harvey, M. Medina-Ramirez, J. D. Roser, R. Smith, D. Westcott, B. F. Keele, J. W. Bess, R. W. Sanders, J. D. Lifson, J. P. Moore, M. Crispin, Site-specific glycosylation of virion-derived HIV-1 Env Is mimicked by a soluble trimeric immunogen. Cell Rep. 24, 1958–1966.e5 (2018).

34. Y. Watanabe, Z. T. Berndsen, J. Raghwani, G. E. Seabright, J. D. Allen, O. G. Pybus, J. S. McLellan, I. A. Wilson, T. A. Bowden, A. B. Ward, M. Crispin, Vulnerabilities in coronavirus glycan shields despite extensive glycosylation. Nat. Commun. 11, 2688 (2020).

35. P. Zhao, J. L. Praissman, O. C. Grant, B. Chen, I. Brief, Y. Cai, T. Xiao, K. E. Rosenbalm, K. Aoki, B. P. Kellman, R. Bridger, D. H. Barouch, M. A. Brindley, N. E. Lewis, M. Tiemeyer, R. J. Woods, L. Wells, Virus-Receptor Interactions of Glycosylated SARS-CoV-2 Spike and Human ACE2 Receptor Combining glycomics-informed glycoproteomics and bioinformatic analyses of variants with molecular dynamics simulations, Zhao et al. detail a role for glycan-protein and glycan-glycan interactions in the SARS-CoV-2 viral Spike protein-ACE2 human receptor complex. Virus-Receptor Interactions of Glycosylated SARS-CoV-2 Spike and Human ACE2 Receptor. Cell Host Microbe. 28, 1–16 (2020).

36. S. J. Morris, S. Sebastian, A. J. Spencer, S. C. Gilbert, Simian adenoviruses as vaccine vectors. Future Virol. 11 (2016), pp. 649–659.

37. M. D. J. Dicks, A. J. Spencer, L. Coughlan, K. Bauza, S. C. Gilbert, A. V. S. Hill, M. G. Cottingham, Differential immunogenicity between HAdV-5 and chimpanzee adenovirus vector ChAdOx1 is independent of fiber and penton RGD loop sequences in mice. Sci. Rep. 5, 1–15 (2015).

38. R. Essalmani, J. Jain, D. Susan-Resiga, U. Andréo, A. Evagelidis, R. M. Derbali, D. N. Huynh, F. Dallaire, M. Laporte, A. Delpal, P. Sutto-Ortiz, B. Coutard, C. Mapa, K. Wilcoxen, É. Decroly, T. N. Pham, É. A. Cohen, N. G. Seidah, bioRxiv, in press, doi:10.1101/2020.12.18.423106.

39. M. Uhlen, L. Fagerberg, B. M. Hallstrom, C. Lindskog, P. Oksvold, A. Mardinoglu, A. Sivertsson, C. Kampf, E. Sjostedt, A. Asplund, I. Olsson, K. Edlund, E. Lundberg, S. Navani, C. A.-K. Szigyarto, J. Odeberg, D. Djureinovic, J. O. Takanen, S. Hober, T. Alm, P.-H. Edqvist, H. Berling, H. Tegel, J. Mulder, J. Rockberg, P. Nilsson, J. M. Schwenk, M. Hamsten, K. von Feilitzen, M. Forsberg, L. Persson, F. Johansson, M. Zwahlen, G. von Heijne, J. Nielsen, F. Ponten, Tissue-based map of the human proteome. Science. 347, 1260419–1260419 (2015).

40. I. Hamming, W. Timens, M. L. C. Bulthuis, A. T. Lely, G. J. Navis, H. van Goor, Tissue distribution of ACE2 protein, the functional receptor for SARS coronavirus. A first step in understanding SARS pathogenesis. J. Pathol. 203, 631–637 (2004).

41. W. Sungnak, N. Huang, C. Bécavin, M. Berg, R. Queen, M. Litvinukova, C. Talavera-López, H. Maatz, D. Reichart, F. Sampaziotis, K. B. Worlock, M. Yoshida, J. L. Barnes, N. E. Banovich, P. Barbry, A. Brazma, J. Collin, T. J. Desai, T. E. Duong, O. Eickelberg, C. Falk, M. Farzan, I. Glass, R. K. Gupta, M. Haniffa, P. Horvath, N. Hubner, D. Hung, N. Kaminski, M. Krasnow, J. A. Kropski, M. Kuhnemund, M. Lako, H. Lee, S. Leroy, S. Linnarson, J. Lundeberg, K. B. Meyer, Z. Miao, A. V. Misharin, M. C. Nawijn, M. Z. Nikolic, M. Noseda, J. Ordovas-Montanes, G. Y. Oudit, D. Pe’er, J. Powell, S. Quake, J. Rajagopal, P. R. Tata, E. L. Rawlins, A. Regev, P. A. Reyfman, O. Rozenblatt-Rosen, K. Saeb-Parsy, C. Samakovlis, H. B. Schiller, J. L. Schultze, M. A. Seibold, C. E. Seidman, J. G. Seidman, A. K. Shalek, D. Shepherd, J. Spence, A. Spira, X. Sun, S. A. Teichmann, F. J. Theis, A. M. Tsankov, L. Vallier, M. van den Berge, J. Whitsett, R. Xavier, Y. Xu, L. E. Zaragosi, D. Zerti, H. Zhang, K. Zhang, M. Rojas, F. Figueiredo, SARS-CoV-2 entry factors are highly expressed in nasal epithelial cells together with innate immune genes. Nat. Med. 26, 681–687 (2020).

42. R. Kornfeld, S. Kornfeld, Assembly of asparagine-linked oligosaccharides. Annu. Rev. Biochem. 54, 631–664 (1985).

43. P. Gagneux, M. Aebi, A. Varki, Evolution of Glycan Diversity (Cold Spring Harbor Laboratory Press, 2015).

44. D. N. Mastronarde, Automated electron microscope tomography using robust prediction of specimen movements. J Struct Biol. 152, 36–51 (2005).

45. W. J. H. Hagen, W. Wan, J. A. G. Briggs, Implementation of a cryo-electron tomography tilt-scheme optimized for high resolution subtomogram averaging. J. Struct. Biol. 197, 191–198 (2017).

46. S. Q. Zheng, E. Palovcak, J. P. Armache, K. A. Verba, Y. Cheng, D. A. Agard, MotionCor2: anisotropic correction of beam-induced motion for improved cryo-electron microscopy. Nat Methods. 14, 331–332 (2017).

47. J. R. Kremer, D. N. Mastronarde, J. R. McIntosh, Computer visualization of three-dimensional image data using IMOD. J Struct Biol. 116, 71–76 (1996).

48. E. F. Pettersen, T. D. Goddard, C. C. Huang, G. S. Couch, D. M. Greenblatt, E. C. Meng, T. E. Ferrin, UCSF Chimera – A visualization system for exploratory research and analysis. J. Comput. Chem. 25, 1605–1612 (2004).

49. A. Shevchenko, H. Tomas, J. Havliš, J. V. Olsen, M. Mann, In-gel digestion for mass spectrometric characterization of proteins and proteomes. Nat. Protoc. 1, 2856–2860 (2007).

50. P. Emsley, M. Crispin, Structural analysis of glycoproteins: Building N-linked glycans with coot. Acta Crystallogr. Sect. D Struct. Biol. 74, 256–263 (2018).

