## Supplementary Information for "Native-like SARS-CoV-2 spike glycoprotein expressed by ChAdOx1 nCoV-19/AZD1222 vaccine"

This document includes:

Supplementary Figures 1-4

Supplementary Table 1

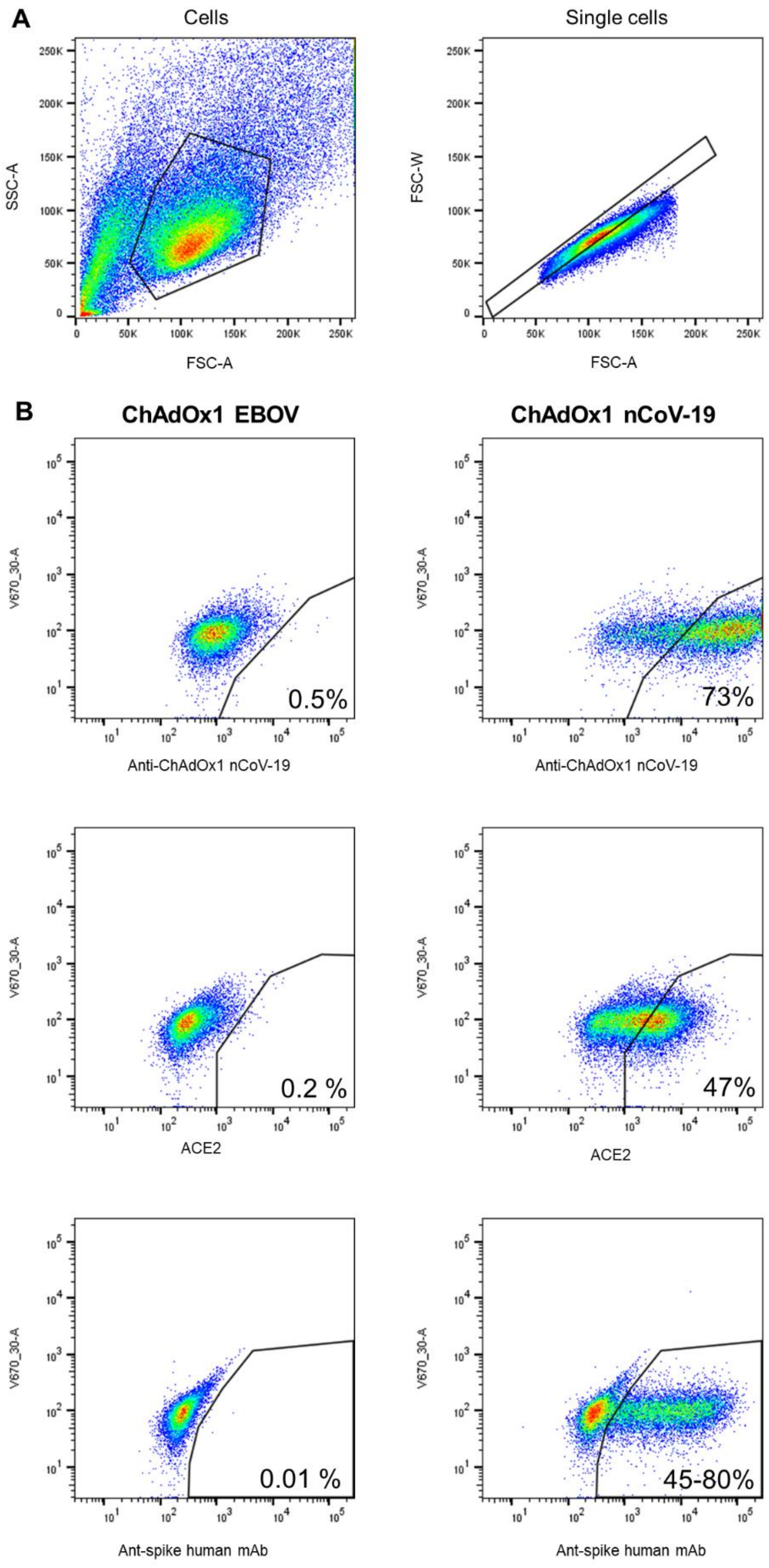


**Supplementary Figure 1. Gating strategy for vaccine expression FACS**

**(A)** Single cells were isolated for further analysis. **(B)** Percentages of positive cells were determined using the gating strategy determined in uninfected cells and displayed for ChAdOx1- EBOV (left panels) and ChAdOx1-nCoV19 (right panels) infected cells, with percentage of positive cells indicated in gate. Representative data is shown, for human mAbs data for mAb 71 is shown.

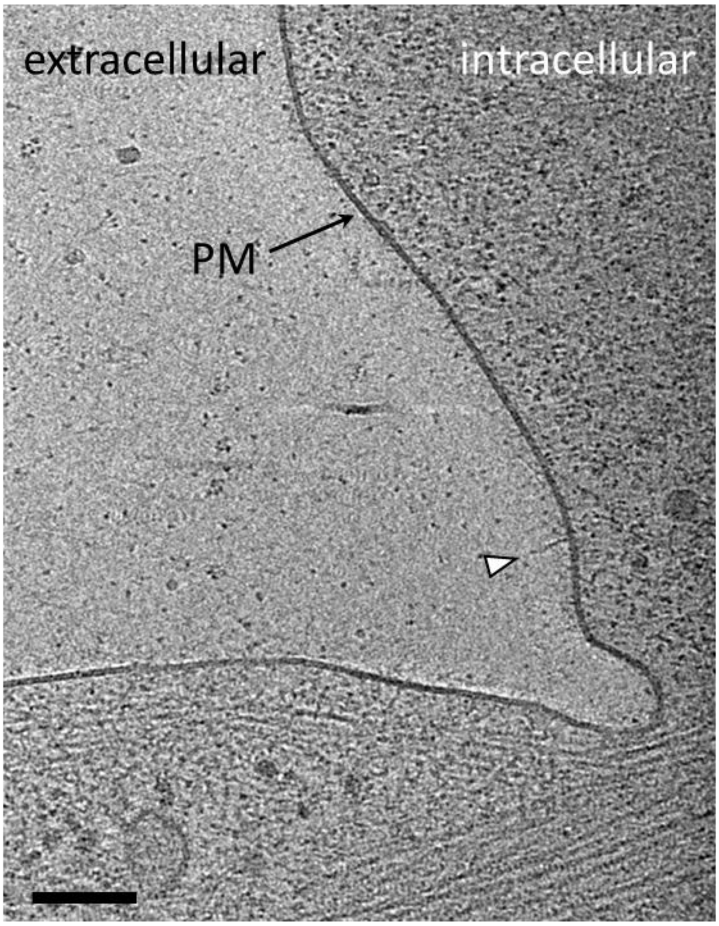


**Supplementary Figure 2. Non-transduced cells lack prefusion-like surface receptors.** Tomographic slice of U2OS control cell surface. PM − Plasma membrane. Slice is 2.13 Å thick. White arrowhead point to a thin and elongated surface receptor. Scale bar is 100 nm.

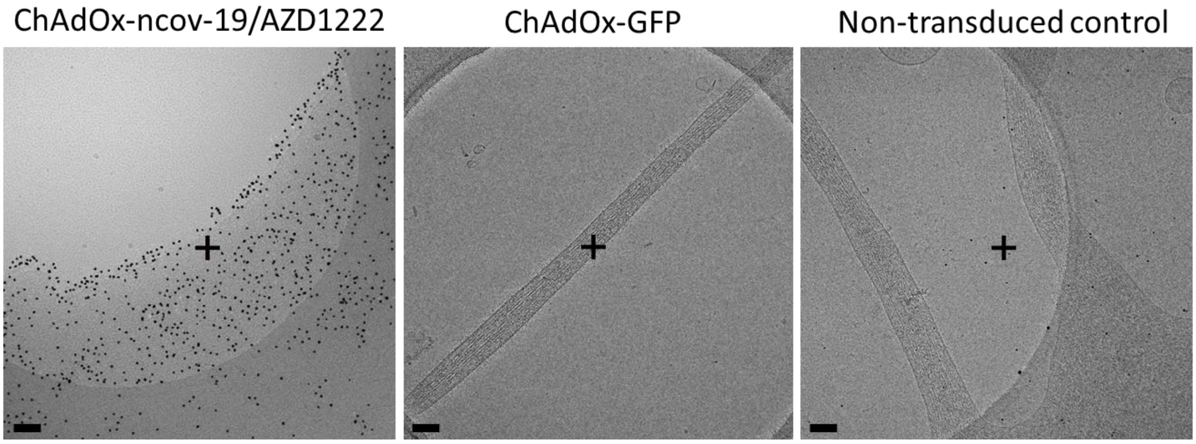


**Supplementary Figure 3. Cryoimmunolabelling of ChAdOx-nCoV-19/AZD1222 derived spike**. Cryo-EM image of U2OS cells transduced with ChAdOx-nCoV-19/AZD1222, ChAdOx-GFP and non-transduced controls. Cells were incubated with ChAdOx-nCoV-19/AZD1222 vaccinated mice sera and labelled with anti-mouse Fab conjugated with 10nm Au beads prior to plunge freezing. Scale bar is 100 nm.

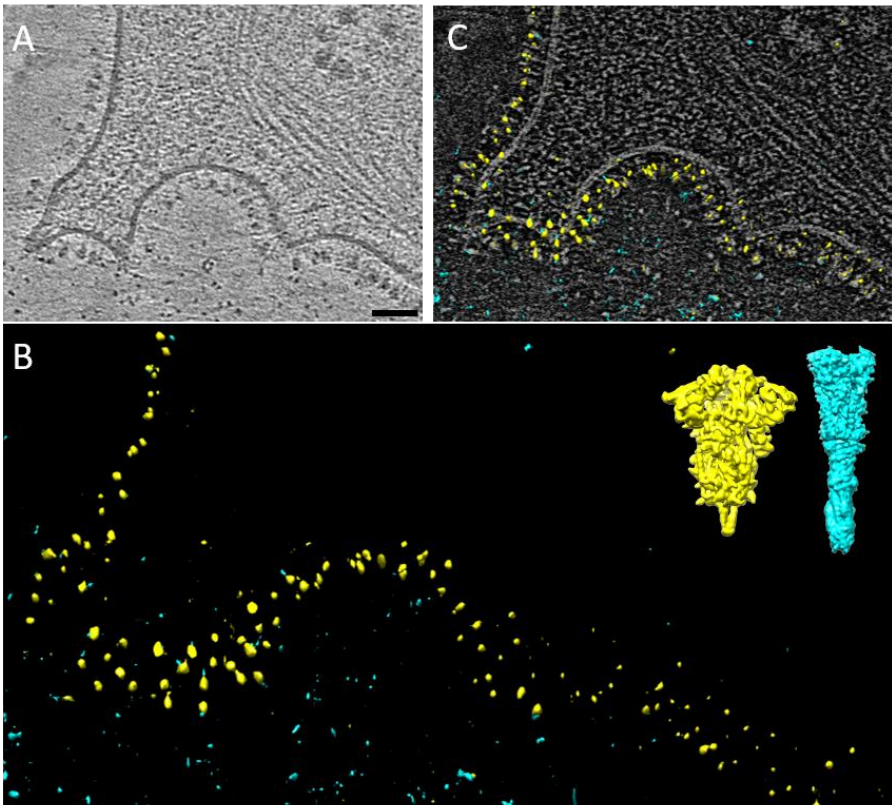


**Supplementary Figure 4.** Template searching for pre- and post-fusion spike proteins. (A) Tomographic slice of a ChAdOx nCoV-19 infected U2OS cell. (B) Results of template matching by cross-correlation of tomographic volume with either a pre-fusion spike reference (EMD-21452, yellow) or a post-fusion spike reference (EMD-7040, cyan). The resulting cross-correlation maps are projected through 18.5 nm thick volume. Inset, the pre- (yellow) and post-fusion (cyan) spike references. (C) The template matching results shown in (B) overlaid with a 1.6 nm thick tomographic slice in (A). Scale bar is 50 nm.

**Supplementary Table 1. Glycoform abundances observed across S0 and S1/S2 SARS-CoV-2 spike protein derived from ChAdOx1 nCoV-19.**

Cleaved

| | N17 | N61 | N74 | N122 | N149 | N165 | N234 | N282 | N331 | N343 | N603 | N616 | N657 | N709 | N717 | N801 | N1074 | N1098 | N1134 | N1158 | N1173 | N1194 | Total |
|---|---|---|---|---|---|---|---|---|---|---|---|---|---|---|---|---|---|---|---|---|---|---|---|
| Mannose/Hybrid | 0% | 81% | 0% | 60% | n/a | 0% | 94% | 18% | 11% | 74% | 0% | 100% | 100% | 100% | 100% | 82% | 100% | 25% | n/a | n/a | n/a | n/a | 56% |
| Complex | 0% | 19% | 100% | 40% | n/a | 100% | 6% | 82% | 89% | 26% | 100% | 0% | 0% | 0% | 0% | 18% | 0% | 75% | n/a | n/a | n/a | n/a | 38% |
| Unoccupied | 100% | 0% | 0% | 0% | n/a | 0% | 0% | 0% | 0% | 0% | 0% | 0% | 0% | 0% | 0% | 0% | 0% | 0% | n/a | n/a | n/a | n/a | 6% |

Uncleaved

| | N17 | N61 | N74 | N122 | N149 | N165 | N234 | N282 | N331 | N343 | N603 | N616 | N657 | N709 | N717 | N801 | N1074 | N1098 | N1134 | N1158 | N1173 | N1194 | Total |
|---|---|---|---|---|---|---|---|---|---|---|---|---|---|---|---|---|---|---|---|---|---|---|---|
| Mannose/Hybrid | n/a | 100% | 100% | 81% | n/a | 72% | 98% | 74% | 78% | 69% | 69% | 100% | 100% | 100% | 100% | 66% | 86% | 69% | 100% | n/a | n/a | 73% | 85% |
| Complex | n/a | 0% | 0% | 19% | n/a | 28% | 2% | 26% | 22% | 31% | 31% | 0% | 0% | 0% | 0% | 34% | 14% | 31% | 0% | n/a | n/a | 27% | 15% |
| Unoccupied | n/a | 0% | 0% | 0% | n/a | 0% | 0% | 0% | 0% | 0% | 0% | 0% | 0% | 0% | 0% | 0% | 0% | 0% | 0% | n/a | n/a | 0% | 0% |
